# Mechanism of ozone alleviation of malignant ascites in hepatocellular carcinoma through inhibition of NETs

**DOI:** 10.1101/2023.04.26.538380

**Authors:** Feng Han, Jiayou Guo, Mingchen Mu, Ka Bian, Zhenting Cui, Qiong Duan, Jianxin Ma, Lai Jin, Wentao Liu, Fanghong Chen

**Affiliations:** Department of Oncology, Xuzhou Medical College,Xuzhou,221004, Jiangsu Province, China; Department of Oncology, Lianyungang oriental Hospital,Lianyungang,222042, Jiangsu Province, China; Department of infectious diseases, Lianyungang oriental Hospital, Lianyungang 222042, Jiangsu Province, China; Department of Oncology, Bengbu Medical College,Bengbu,233030,Anhui Province,China; Institute of Translational Medicine, Nanjing Medical University, Nanjing, P.R.211166, Jiangsu Province,China; Department of Center for Clinical Research and Translational Medical, Lianyungang oriental Hospital,Lianyungang,222042, Jiangsu Province, China

**Keywords:** neutrophil extracellular trap network, Ozone, hepatocellular carcinoma ascites

## Abstract

Malignant ascites in hepatocellular carcinoma is usually a sign of advanced disease and poor prognosis and is also thought to be associated with chronic inflammation mediated by neutrophil extracellular trap networks (NETs). Whereas ozone, a strong oxidant, has significant antibacterial and anti-inflammatory effects, its effectiveness in treating malignant liver ascites is unclear.**c**Ozone showed significant anti-peritoneal fluid production properties in H22 tumor-bearing mice, and ozone reduced peritoneal fluid production by activating AMPK and up-regulating SR-A phagocytosis damage-associated molecular patterns (DAMPs) to reduce the production of NETs. This suggests that ozone could be used as a new drug for the treatment of malignant ascites in hepatocellular carcinoma.

## 1. Introduction

Malignant ascites (MA) is a common complication of late-stage tumors such as liver, ovarian, pancreatic and gastric cancers [1,2]. The most common type of malignant ascites in hepatocellular carcinoma is characterized by persistence, high volume and recurrent occurrence. Due to increased intra-abdominal pressure and pain, patients have a poor quality of survival, exhibiting nausea, vomit and anorexia, and severe ascites can even be life-threatening. Thus, it is an urgent clinical problem caused by cancer to be solved. Currently, chemotherapy, laparotomy, diuretics and abdominal venous shunts are widely used to relieve the symptoms of hepatocellular carcinoma ascites [3,4]. However, these methods only relieve symptoms but do not address the underlying problem, and have many side effects [5,6]. Therefore, to search for more effective treatments for malignant ascites in hepatocellular carcinoma has become a major focus of current drug research.

The significant increase of inflammatory cytokines, often accompanied by an increase of neutrophils in the ascites, is a hallmark of tumor progression[7]. In advanced stages of cancer, apoptotic necrosis releases a large numbers of DAMPs, which activate TLR4 receptors and eventually release nuclear contents from neutrophils, resulting in the formation of NETs, a new mode of neutrophil defense and death distinct from apoptosis and necrosis [8,9,10]. NETs play an important role in the body’s resistance to infection, but also induce inflammatory storms that directly or indirectly damage cells and tissues, increase vascular permeability, and promote tumor growth and metastasis [11,12,13,14]. Therefore, it is essential to reduce the production of NETs for the remission of MA.

There is some evidence that ozone treatment, widely used in traditional medicine, has antibacterial properties, stimulates oxygen metabolism and activates the immune system, and that medical ozone is clinically effective in the treatment of herniated discs [15], joint pain [16], diabetic ulcers [17] and chronic ulcerative colitis [18].

Furthermore, our previous studies have shown that ozone treatment enhanced SR-A phagocytosis of high mobility group protein-1 (HMGB1), a core component of NETs, by activating AMPK, which is also recognized by inflammatory cells as DAMPs and stimulates inflammatory responses [19]. In other words, ozone treatment may be an effective therapy for malignant ascites as it reduces both the NETs activation by phagocytosis of DAMPs and the NETs formation.

The aim of this study was to investigate the possible anti-ascites properties of ozone. In addition, a mouse ascites xenograft model was developed to investigate the potential mechanisms of the anti-ascites effect of ozone in vivo.

## 2. Materials and Methods

### 2.1 Cell culture

The H22cell line was obtained from the Shanghai Institute of Cell Biology (Shanghai, China). They were cultured in DMEM medium (Gibco, Grand Island, NY, USA), supplemented with 10% fetal bovine serum (FBS, Hyclone, Logan, UT, USA), and 1% penicillin/streptomycin (Invitrogen Inc., Carlsbad, CA, USA) (100 units/mL). Cells were kept under of 5% CO2 in an incubator at 37°C.

### 2.2 Clinical data

There were 10 patients with stage III-VI primary liver cancer, including 6 males and 4 females, aged 45-75 years, with a median of 63 years. All were inpatients of our hospital from May 2022 to December 2022. All cases were diagnosed as primary hepatocellular carcinoma. 10 healthy control subjects were normal subjects who were examined in our hospital. Fatty liver, liver cirrhosis and hepatitis were excluded. We had access to information that could identify individual participants during or after data collection.All patients in this study signed an informed consent form before the operation and were approved by our ethics committee (approval number: 2022-036-01).

### 2.3 Animals and Tumor Formation

Male ICR mice (20-22g, 4–6weeks old) were provided by Shanghai Biocay Biotechnology Co., Ltd., China(All animal experiments were approved by Nanjing Medical University Animal Care and Use Committee and were approved by the Ethics Committee of Nanjing Medical University (No. IACUC-1908026). Animals were housed under standard specific pathogen-free (SPF) conditions (25° C±1° C and 55%±5% relative humidity). Mice were fed a standard pelleted diet and provided with water ad libitum. These animals were acclimatised and fed for seven days prior to the experiment. The experimental mice were handled in accordance with animal ethical requirements and were approved by the ethics committee of our institution. H22 cells at logarithmic growth stage were taken and resuspended in saline at a concentration of 1×107 cells/ml. 36 ICR mice were randomly selected to receive 0.2mL of H22 cell suspension (2×106 cells per mouse) in the left abdominal cavity, with the remaining 12 being a blank group. H22 tumour-bearing mice were randomly divided into three groups (n=12 per group): model group; ozone-treated group (15ug/ml and 30ug/ml). The ozone treatment group was treated with continuous intraperitoneal injection of 0.5 ml per mouse for thirteen days. Ozone is produced by a medical ozone therapy device (Ozomed Smart, kastner, Germany).

Ozone obtained from medicinal grade oxygen was used immediately. The abdominal girth and body weight of each mouse were measured daily. Half of the mice in each group were executed on day thirteen and the small intestinal tissue and ascites were removed, the rest of the mice were fed normally; their survival time was recorded.

### 2.4 cf-DNA measurement

The QUANT-iT PicoGreenTMds DNA AssayKit (Invitrogen Inc., USA) was used for the detection of cf-DNA, first make PicoGreen working solution by diluting PicoGreen storage solution 200 times with 1 x TE buffer and store away from light. Collect 5 ml of peripheral blood from healthy human and hepatocellular carcinoma patients or ascites from mice, centrifuge at 3000 rpm for 15 min and collect the serum or ascites supernatant. Add 50 μl of serum or ascites supernatant to the luminescent labeling plate, followed by an equal amount of PicoGreen® dsDNA Reagent working solution and carefully shake and mix; the reaction is carried out at room temperature and protected from light for 5 min; the fluorescence intensity is measured at 480/520 nm by luminescent labeling instrument (SynergyH1,BioTek,USA).

### 2.5 Immunofluorescence

Mouse intestinal tissue was fixed in 4% formaldehyde, paraffin-embedded and sectioned (5 μm). All immunofluorescence steps were carried out according to the Cell Signaling Technology immunofluorescence protocol. Specifically, antigen retrieval was performed using Sodium Citrate buffer pH 6 for all slides. Rabbit anti-CitH3 polyclonal antibody (1:200, ab5103, Abcam) and mouse anti-MPO monoclonal antibody (1:200, ab90811, Abcam) were added and incubated overnight, then incubated with goat anti-rabbit IgG Alexa Fluor 647 polyclonal antibody (A21245, Invitrogen) and rabbit anti-mouse IgG Alexa Fluor 488 polyclonal antibody (A21245, Invitrogen) for 90min at room temperature. DNA was counterstained with 10 µM DAPI (MBD0015, Sigma-Aldrich, USA). Final photography and analysis were finished by a fully automated pathology scanner (Pannoramic 250 FLASH, 3D HISTECH, Hungary).

### 2.6 Ascites Cytokine Analyses

The ascites fluid was extracted and centrifuged, and the supernatant was collected. The levels of cytokine, including IL-6, IFN-γ, TNF-α, MMP-9and VEGF, were measured by commercial ELISA kits (Neobioscience, Shenzhen, China) according to the manufacturer’s instructions.

### 2.7 Immunocytochemistry

Mouse intestinal tissues were fixed in 4% buffered formaldehyde, embedded in paraffifin, and cut as 5 mm thick slides. Antigen retrieval was performed by heating for 3 min in 10 mM sodium citrate buffer (pH 6.0). The slides were immersed in 3% hydrogen peroxide for 20 min to block the activity of endogenous peroxidase, then incubated overnight at 4°C with an anti-AMPK phosphorylation(Thr172, AF5908, Beyotime, Shanghai, China) or anti-SR-A (MSR1, 17858-1-Ap, Proteinrech, Chicago, IL, USA) polyclonal antibodies at a 1:200 dilution and incubated with biotinylated goat anti-rabbit antibodies for 1 h at room temperature (PA2202, Proteinbio, Nanjing, China). Then, the slides were washed with PBS and additionated with diaminobenzidine (DAB) chromogen for 3–5 min to yield a dark brown color. The slides were counterstained with hematoxylin and analyzed under a light microscope (Olympus, Tokyo, Japan).

### 2.8 Western Blot Analysis of P-AMPK and SR-A

The samples of ascites fluid sediment of mice were collected and lysed using in radio immunoprecipitation assay (RIPA) lysis buffer (Beyotime, Shanghai, China) supplemented with protease inhibitors (1% PMSF, Beyotime, Shanghai, China). Equal amounts of proteins were separated in 8% SDS/PAGE gels and transferred to polyvinylidene diflfluoride (PVDF) membranes. The membranes were blocked with 0.5% skim milk dissolved in 1 xTBST (Tris buffered saline with Tween 20; pH 7.5) at room temperature for 30 min, incubated with rabbit anti-P-AMPK (1:1000), rabbit anti-SR-A (1:1000), or rabbit anti-β-actin (1:2000; FB1033, Proteinbio, Nanjing, China) overnight at 4°C, and incubated with goat anti-rabbit IgG-HRP (1:4000) for 1 h at room temperature. Finally, the immunoreactive bands were detected by enhanced chemiluminescence. The target bands were analyzed using Image Lab software.

### 2.9 Statistical analysis

Graphpad prism9 (GraphPad9, San Diego, CA, USA)statistical software was used to analyze and create graphs. data were expressed as mean ± standard deviation, t-test was used for comparison between two groups and one-way ANOVA was used for comparison between more than two groups. p < 0.05 was statistically significant.

## 3. Results

### 3.1 NETs are higher in liver cancer patients compared that in normal subjects

NETs are DNA-based and cf-DNA can be used to indirectly reflect the level of NETs in the body. Peripheral blood was collected from patients with clinical liver cancer, serum was separated, in which dsDNA was stained by PicoGreen, and the samples were detected with 520/480 nm. The results are shown in Figure 1. Compared with the control group, the serum dsDNA levels of patients with liver cancer were increased, and the difference was statistically significant, P<0.01.

**Figure 1.**
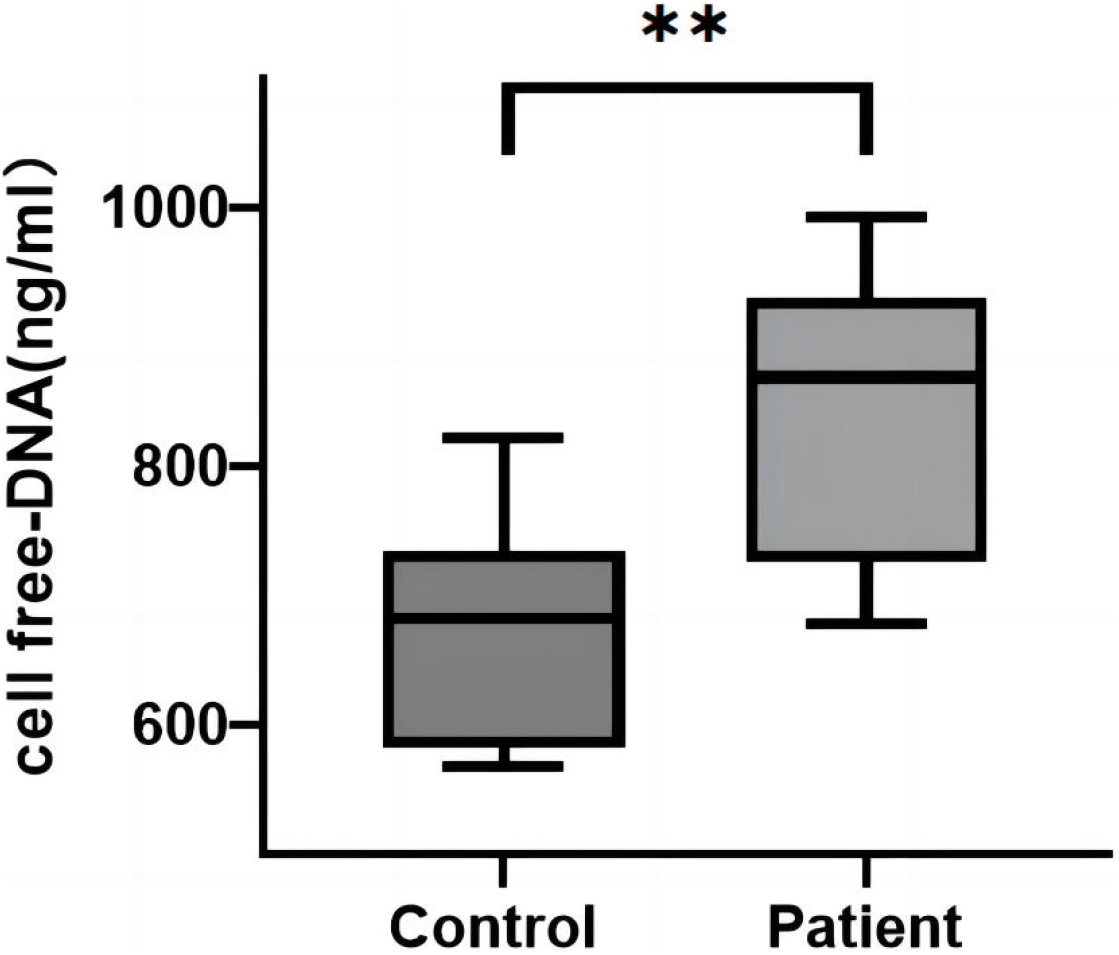
Comparison of free DNA levels in plasma of liver cancer patients and normal subjects. The data are shown as mean ± SD (n = 10). (** p < 0.01).

### 3.2 Anti-ascites effect of ozone therapy in vivo

To investigate whether ozone therapy inhibits the production of hepatocellular carcinoma ascites in vivo, we established a malignant ascites model in ICR mice (Figure 2A). We transplanted H22 cells into the left abdominal cavity of the mice. Seven days later, mice inoculated with tumor cells developed abdominal bulges. The tumour-bearing mice were divided into three groups (n=12) according to the mean body weight: the model group; the 15ug/ml ozone group; and the 30ug/ml ozone group, which were treated with ozone for 13 consecutive days. Eleven days after inoculation with tumor cells, the mice in the model group showed an increase in ascites, a rapid increase in body weight and abdominal girth, difficulty in drinking and eating, laziness, lack of lustre, easy loss of hair, and the phenomenon of vertical hair was obvious. Sixteen days after inoculation with tumor cells, the abdomen of the mice in the model group bulged significantly, and the mice showed apathy and almost no activity and started to die. During the periods of seventh to nineteenth day of tumor cell inoculation, we measured the abdominal girth and body weight of the mice. The abdominal girth and body weight were significantly lower in the ozone treated group compared to that in the model group (Figure 2B, C). Nineteen days after tumor cell inoculation, the abdominal girth and body weight of mice in the ozone treated group were significantly lower than those in the model group (Tables 1 and 2). At the end phase of the ozone treatment, half of the mice in each group were executed, their ascites was extracted, photographed and the volume of ascites was measured. The mice of model group had obvious bloody ascites, while the ozone treated group was significantly lighter in color, and the volume of ascites in the ozone treated group was significantly reduced compared to the model group (Figure 2E, F). Furthermore, as shown in Figure 2D, Table 3, the survival rate and survival time (by the end of day 30) increased in a dose-dependent manner in the ozone treated group. In conclusion, these results suggest that ozone therapy has significant function of anti-ascites production in our H22 liver cancer model.

**Table 1:** Mean body weight of mice in each group.

| Group (n=6) | Weight(g) |  |  |  |
| --- | --- | --- | --- | --- |
|  | Day7 | Day11 | Day15 | Day19 |
| **Control** | 25.50 ± 1.61 | 27.89 ± 1.56 | 30.52 ± 1.41 | 32.90 ± 1.72 |
| **Model** | 26.00 ± 1.60 | 30.52 ± 1.78 | 34.45 ± 1.30 | 37.23 ± 1.65 |
| **ozone (15ug/ml)** | 26.10 ± 1.48 | 30.20 ± 1.18 | 32.95 ± 1.08 | 35.72 ± 1.27* |
| **ozone (30ug/ml)** | 26.00 ± 1.52 | 29.77 ± 1.78 | 31.97 ± 1.60 | 34.30 ± 1.51* |

Compared with the model group (\* $p < 0.05$ ).

**Table 2:** Mean abdominal circumference of mice in each group.

| Group (n=6) | Abdominal(cm) |  |  |  |
| --- | --- | --- | --- | --- |
|  | Day7 | Day11 | Day15 | Day19 |
| **Control** | 7.57 ± 0.61 | 8.05 ± 0.60 | 8.50 ± 0.58 | 8.90 ± 0.72 |
| **Model** | 8.50 ± 0.60 | 9.98 ± 0.67 | 11.2 ± 0.68 | 12.67 ± 0.85 |
| **Ozone (15ug/ml)** | 8.60 ± 0.58 | 9.56 ± 0.53 | 10.5 ± 0.60 | 11.70 ± 0.87* |
| **Ozone (30ug/ml)** | 8.40 ± 0.62 | 9.27 ± 0.41 | 9.80 ± 0.56 | 10.80 ± 0.91* |

Compared with the model group (\* $p < 0.05$ ).

**Table 3:**
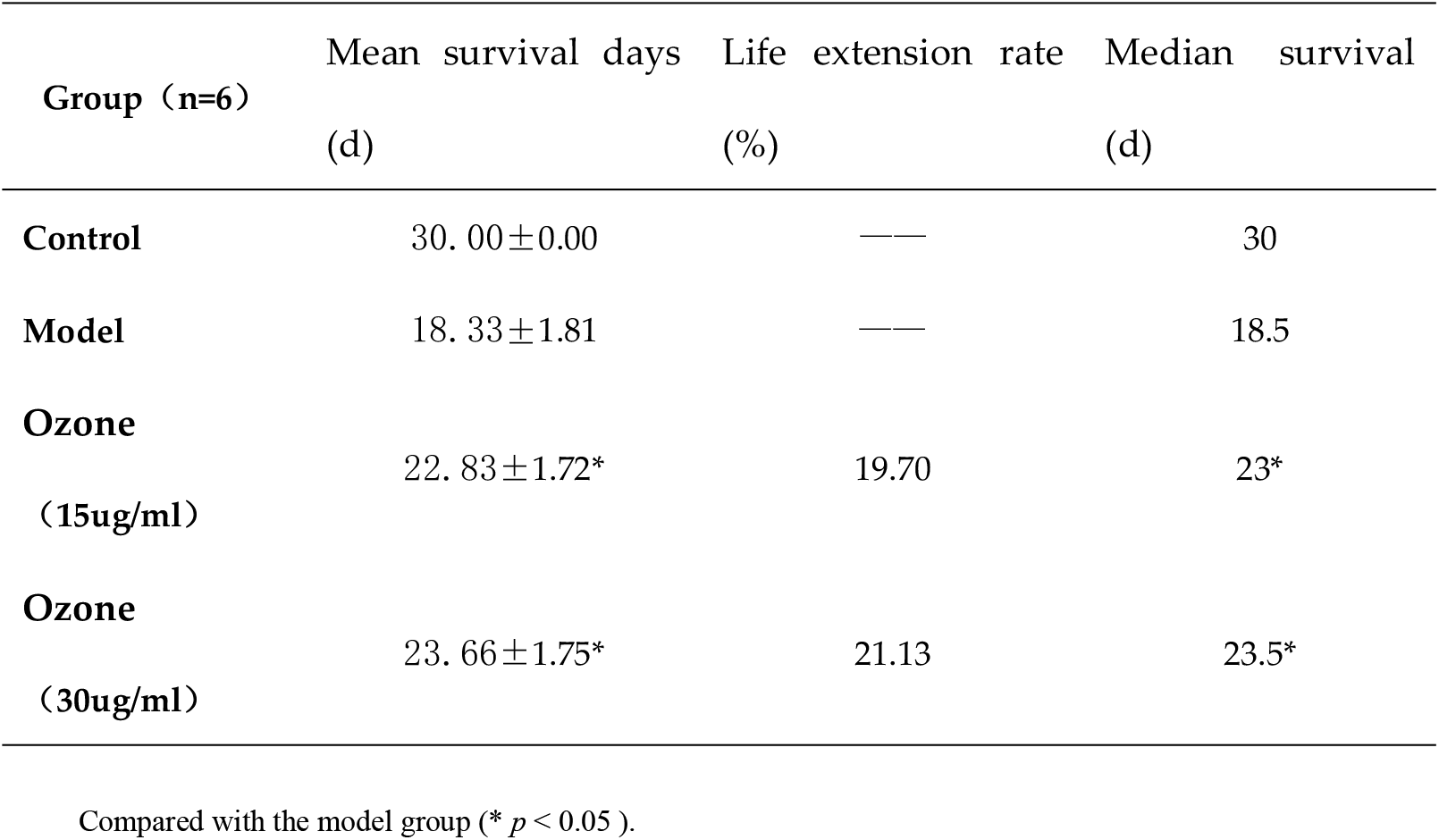
Survival rate of mice in each group.

**Figure 2:**
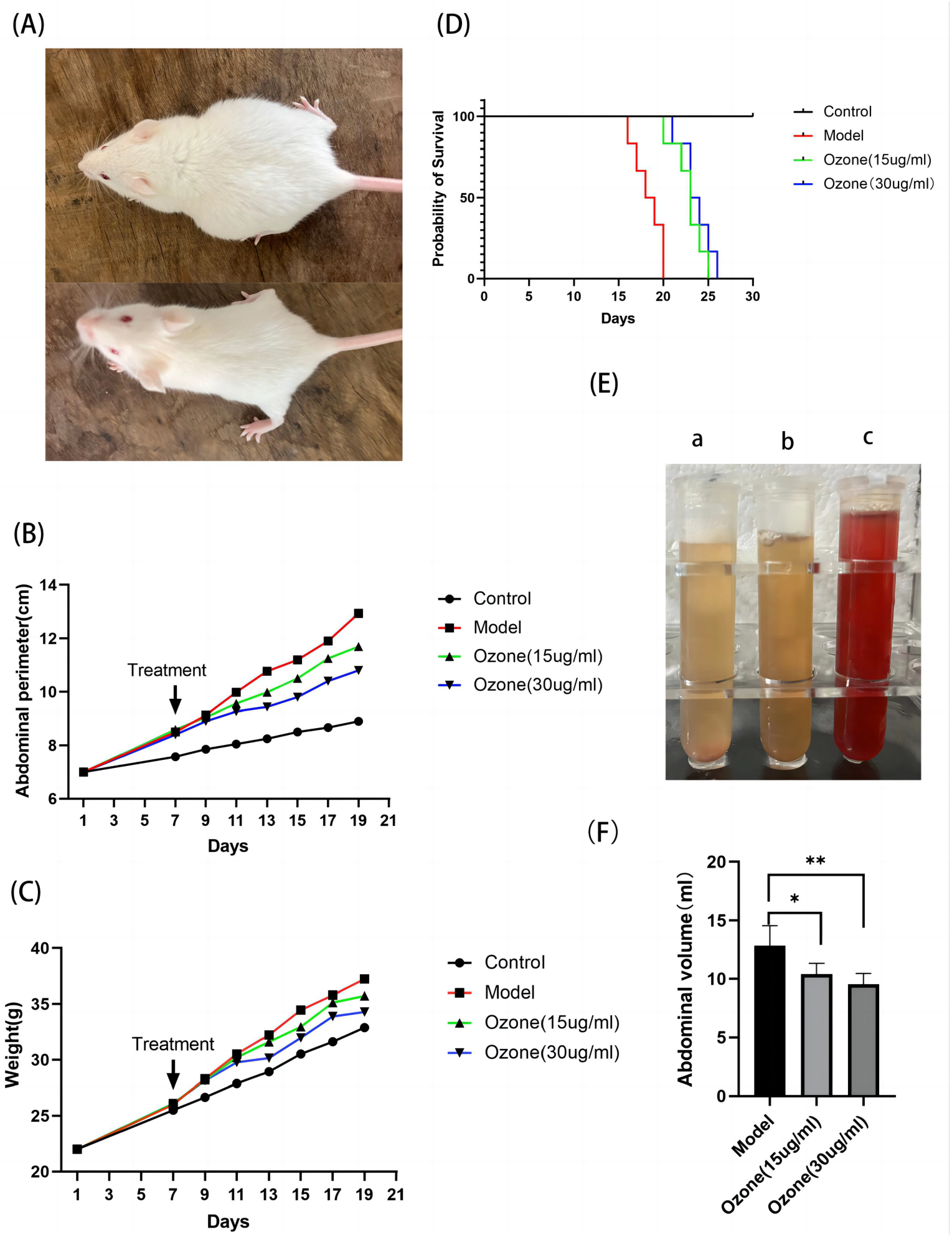
The effect of ozone treatment on H22 liver cancer mice. (A) Mouse model of H22 malignant ascites. (B) Abdominal girth of mice in each group. (C) Mean body weight of mice in each group. (D) Survival curves of mice in each group. (E)Data are expressed as mean ± SD (n=6). (*p<0.05).

### 3.3 Expression and localization of NETs in intestinal tissues of mice with hepatocellular carcinoma ascites

MPO and CitH3 are important components of NETs, which indirectly demonstrate the presence of NETs. We double-stained MPO and CitH3 in mouse intestinal tissues to localize and quantify the expression of NETs. The immunofluorescence results (Figure 3A, B) showed that MPO and CitH3 were barely expressed at the neutrophil aggregates in the intestinal tissues of mice in the control group, while MPO and CitH3 were abundantly overlapped at the neutrophil aggregates of mice in the model group, while fluorescence intensity of MPO and CitH3 was significantly reduced in the ozone-treated group, suggesting the lower level of NETs.

**Figure 3:**
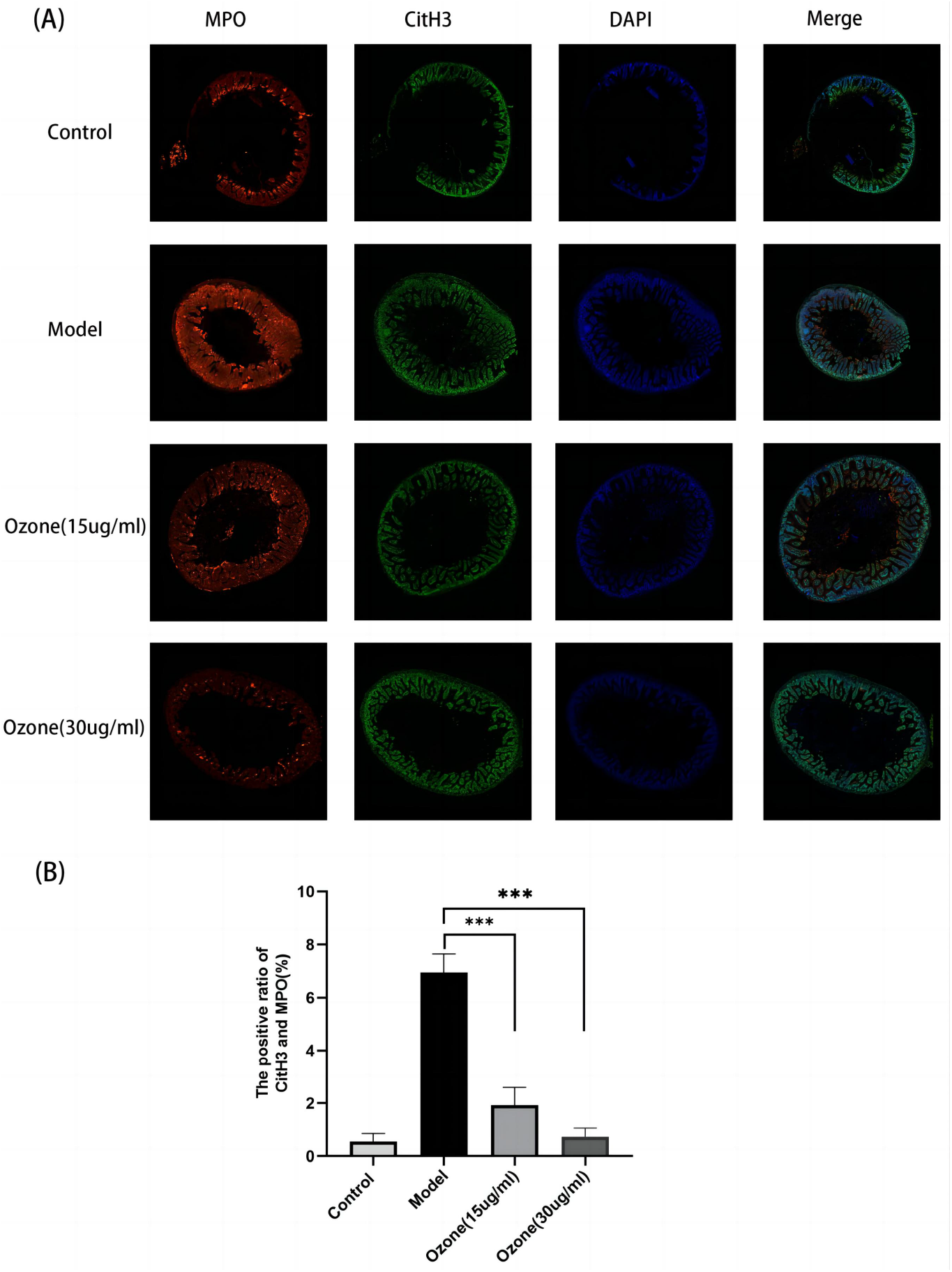
Localization and quantitative analysis of NETs expression in intestinal tissues of various groups of mice. (A).Expression and localization of NETs in each group of mice. (B). Percentage of cells co-expressing CitH3 and MPO.The data are presented as the mean ± SD (n = 6); Compared with the model group (*** p < 0.001).

### 3.4 Effect of ozone on the levels of NETs and cytokines in ascitic fluid

NETs can cause a severe inflammatory response, which is inextricably linked to the development of ascites. We next detected the levels of cf-DNA, IFN-γ, TNF-a, IL-6, VEGF and MMP-9 in ascites by ELISA assay, as shown in Figure 4. The levels of cf-DNA, IFN-γ, TNF-a, IL-6, VEGF and MMP-9 in the ascites of the ozone-treated mice were reduced in a dose-dependent manner compared to those of the model mice.

**Figure 4.**
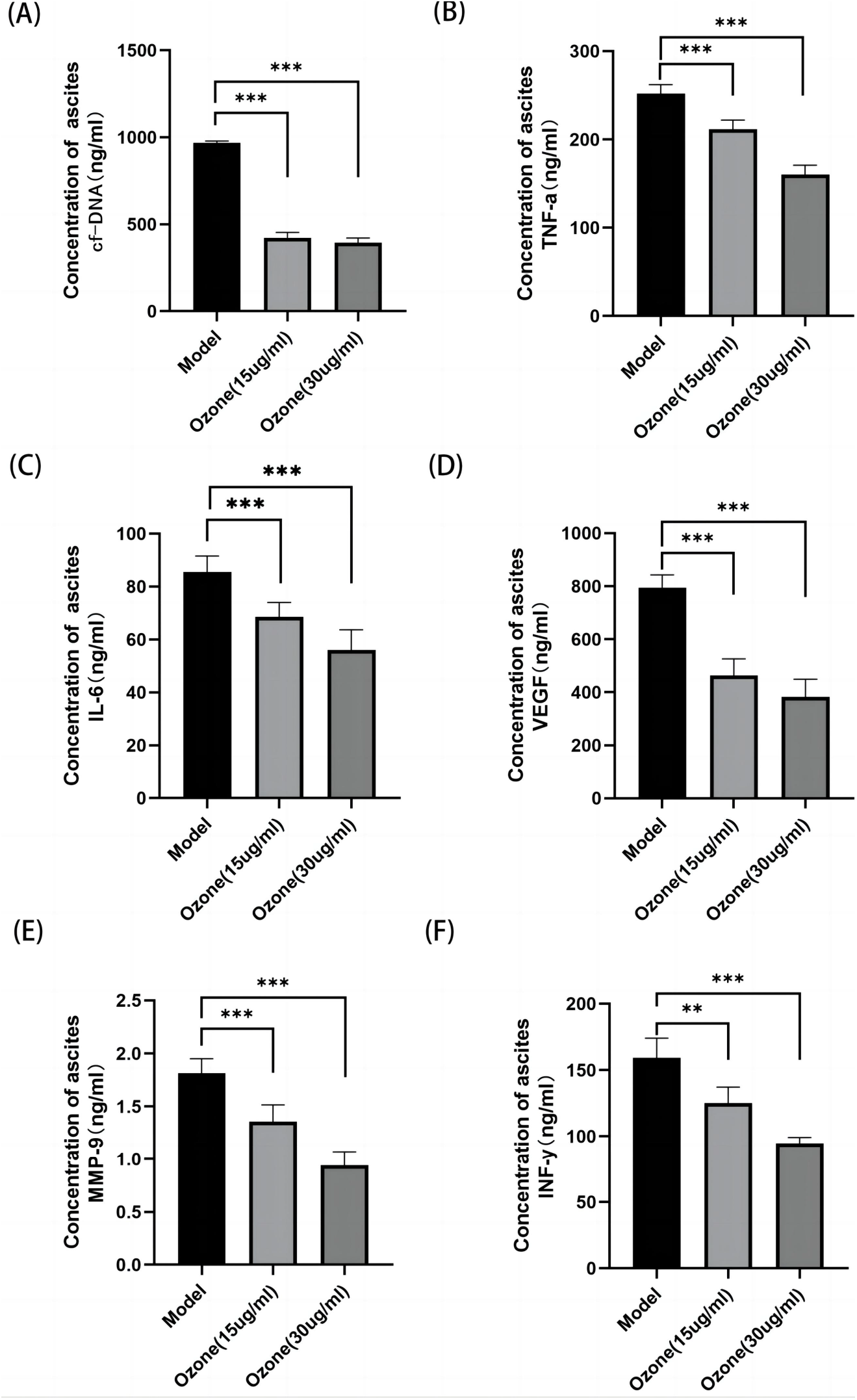
Effect of ozone on NETs and cytokines in the ascites of mice with liver cancer ascites. A cf-DNA, B TNF-a, C IL-6,D VEGF,E MMP-9,F INF-y.The data are presented as the mean ± SD (n = 6); Compared with the model group (** p < 0.01,*** p < 0.001).

### 3.5. Expression of P-AMPK and SR-A in various groups of mice

Our previous studies have shown that the anti-inflammatory effect of ozone is associated with its activation of AMPK, and that SR-A, a downstream factor of AMPK, can also act as an anti-inflammatory agent by enhancing the phagocytosis of macrophages[19,20].To further determine the potential mechanism of which ozone inhibits the production of NETs, we measured the expression of P-AMPK (Thr172) and SR-A proteins in the intestinal mucosa of mice by immunohistochemistry.

P-AMPK and SR-A protein levels were significantly reduced in the model group, which were reversed by ozone therapy. We further examined the expression of P-AMPK and SR-A proteins in ascites sediment. P-AMPK and SR-A protein levels were significantly increased in the ozone-treated group compared to that in the model group (e.g. Figure 5 A). Quantitative analysis showed that ozone significantly activated the AMPK-SR-A pathway (e.g. Fig. 5C). In conclusion, these results suggest that ozone activated AMPK, thereby upregulating SR-A de-phagocytosis of NETs and ultimately reducing the production of malignant ascites.

**Figure 5.**
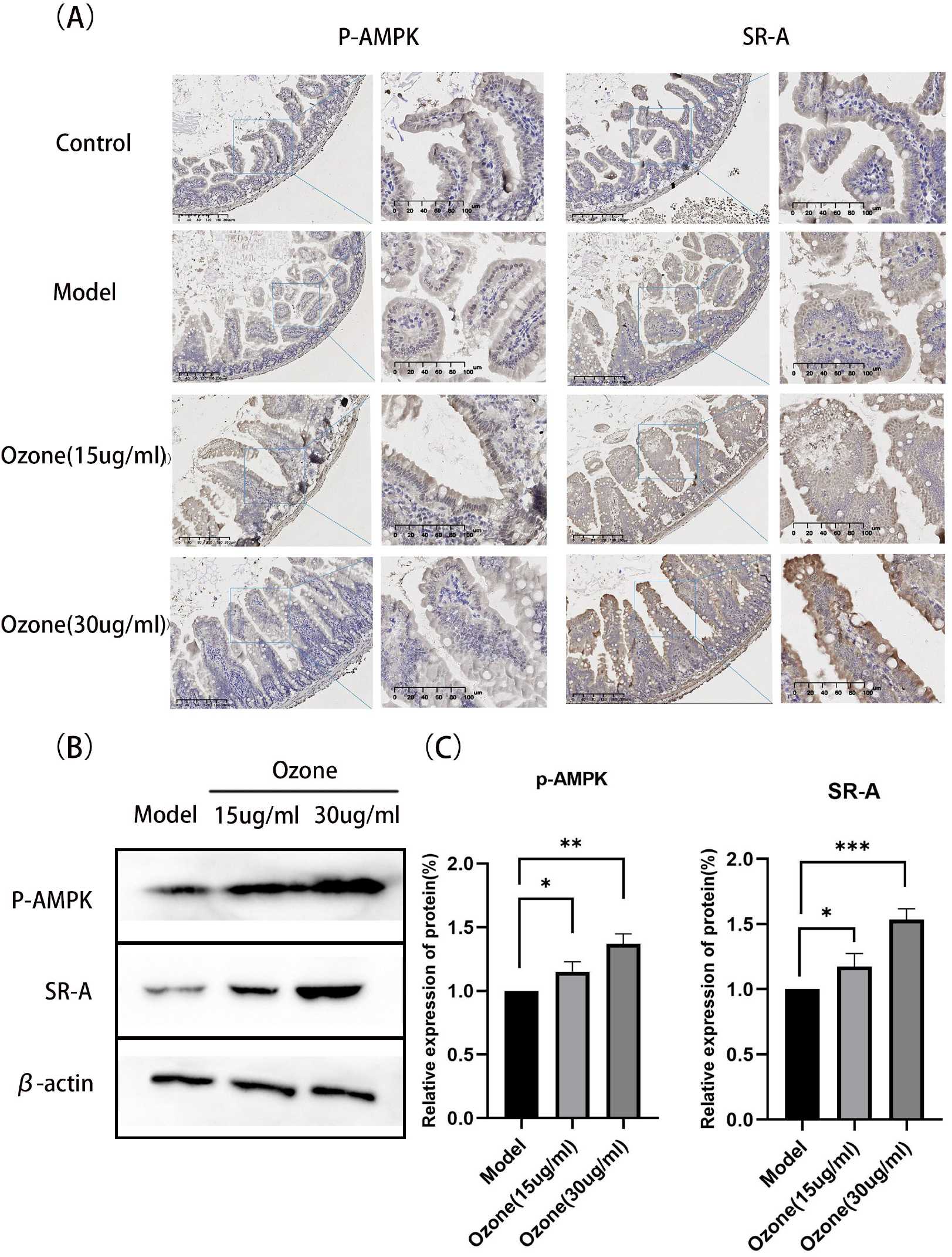
Effect of ozone treatment on p-AMPK and SR-A protein expression. (A) Immunocytochemical images of p-AMPK and SR-A in the intestinal tissues of mice in each group. Moderate and strong brown cytoplasmic staining as positive expression,100×, 200×. (B) Western blot analysis of p-AMPK and SR-A expression (C) Quantitative analysis of relative protein expression in each group. Data are shown as mean ± standard deviation; (*p<0.05, **p<0.01, ***p<0.01) significantly different from the model group.

## 5. Conclusions

In conclusion, our study shows that ozone treatment exhibits significant anti-peritoneal fluid production in a mouse model of ascites. Ozone therapy reduced the formation of NETs at the source, thereby reducing vascular leakage. The present study provides insight into gas therapy for malignant ascites in hepatocellular carcinoma, and further studies are needed to fully elucidate the potential molecular mechanisms underlying the role of ozone in human malignant ascites.

## Disclosure of Ethical Statements

### Approval of the research protocol

The study complied with the Declaration of Helsinki and was approved by the Medical Ethics Committee of Lianyungang Oriental Hospital, Jiangsu Province (2022-036-01).

### Informed Consent

Patient consent was waived due to the retrospective design of this study.

### Registry and the Registration No. of the study/trial

N/A

### Animal Studies

All procedures were strictly performed in accordance with the regulations of the ethics committee of the International Association for the Study of Pain and the Guide for the Care and Use of Laboratory Animals (The Ministry of Science and Technology of China, 2006). All animal experiments were approved by Nanjing Medical University Animal Care and Use Committee and were approved by the Ethics Committee of Nanjing Medical University (No. IACUC-1908026).

### Research involving recombinant DNA

N/A

## Acknowledgments

I am grateful to Professors Ma Jianxin, Jinlai and Liu Wentao for their encouragement and guidance.

## Funding

This research received no external funding.

### Author Contributions

Conceptualization, Lai Jin and Wen Tao Liu; Data curation, Feng Han, Ka Bian, Qiong Duan and Jia Guo; Formal analysis, Feng Han and Ming Mu; Investigation, Feng Han, Ming Mu and Ka Bian; Project administration, Jian Ma; Software, Zhen Cui; Supervision, Jian Ma, Lai Jin and Wen Tao Liu; Validation, Feng Han; Writing – original draft, Feng Han.

### Data Availability Statement

The data presented in this study are available on request from the corresponding author. The data are not publicly available due to medical data privacy issues.

